# Closed-loop sensory feedback enables fast and reliable instrumental acquisition in head-fixed mice

**DOI:** 10.64898/2026.07.26.740810

**Authors:** Ajit Ranganath, Daniel Hähnke, Simon N. Jacob

## Abstract

Instrumental learning typically requires hundreds to thousands of trials in which subjects learn to link motor responses to sensory cues. In standard rodent protocols, response accuracy is reported only at trial end, preventing subjects from correcting erroneously initiated responses. We hypothesized that within-trial, closed-loop sensory feedback would accelerate instrumental learning by providing real-time information about response correctness. Head-fixed mice performed a two-alternative forced-choice task by rotating a choice wheel in response to sensory cues. Mice received either no feedback (n = 18), auditory feedback (n = 16), or audiovisual feedback (n = 4) coupled to wheel movements. Feedback-receiving mice required significantly fewer trials to reach 70 % accuracy criterion (median: 3186, 4918 and 7329 trials for multimodal, unimodal and no feedback, respectively; p = 0.0245) and showed higher accuracy when modifying choices (expert stage: 17 %, 11 % and 9 % accuracy in trials with modified choices for multimodal, unimodal and no feedback, respectively; p=1.04×10⁻⁶). Only feedback mice displayed movement refinements across training (p = 8.02×10⁻⁶, p = 4.07×10⁻¹⁸ and p = 0.2602 for multimodal, unimodal and no feedback, respectively). In summary, closed-loop sensory feedback accelerated instrumental acquisition, demonstrating its value as routine training protocol.

**HIGHLIGHTS:**

- Mice provided with feedback require fewer trials to reach expert stage in an instrumental learning task
- Mice provided with feedback perform with higher accuracy in trials involving ‘changes of mind’
- Movement trajectories of mice provided with feedback undergo refinement as training advances
- Sensory feedback can be used as a training aid to accelerate instrumental learning

## INTRODUCTION

A central goal in neuroscience is to systematically link brain dynamics with behavior ^1,2^. Unraveling the neural circuit mechanisms responsible for intelligent behaviors requires experimental access to neuronal populations for measurements and manipulations ^3^. Rodents, mice in particular, have emerged as an unparalleled model species to study perception and cognition ^4,5^ due to their behavioral tractability, availability of established inbred lines ^6^, transgenic access ^7–9^ and availability of brain atlases ^10–12^. Mice have been used in the freely moving or the head-fixed preparation, depending on the experimental questions at hand. In particular, the head-fixed preparation (**Fig. 1A**) allows for spatially and temporally precise stimulus delivery and has enabled the use of mice in studies of sensory discrimination behaviors involving visual ^13,14^, auditory ^15,16^, tactile ^17^ and odorant ^18–20^ stimuli. Since sensory discrimination is often a steppingstone towards sophisticated cognitive behaviors, the head fixed preparation is well suited in this regard.

**Figure 1:**
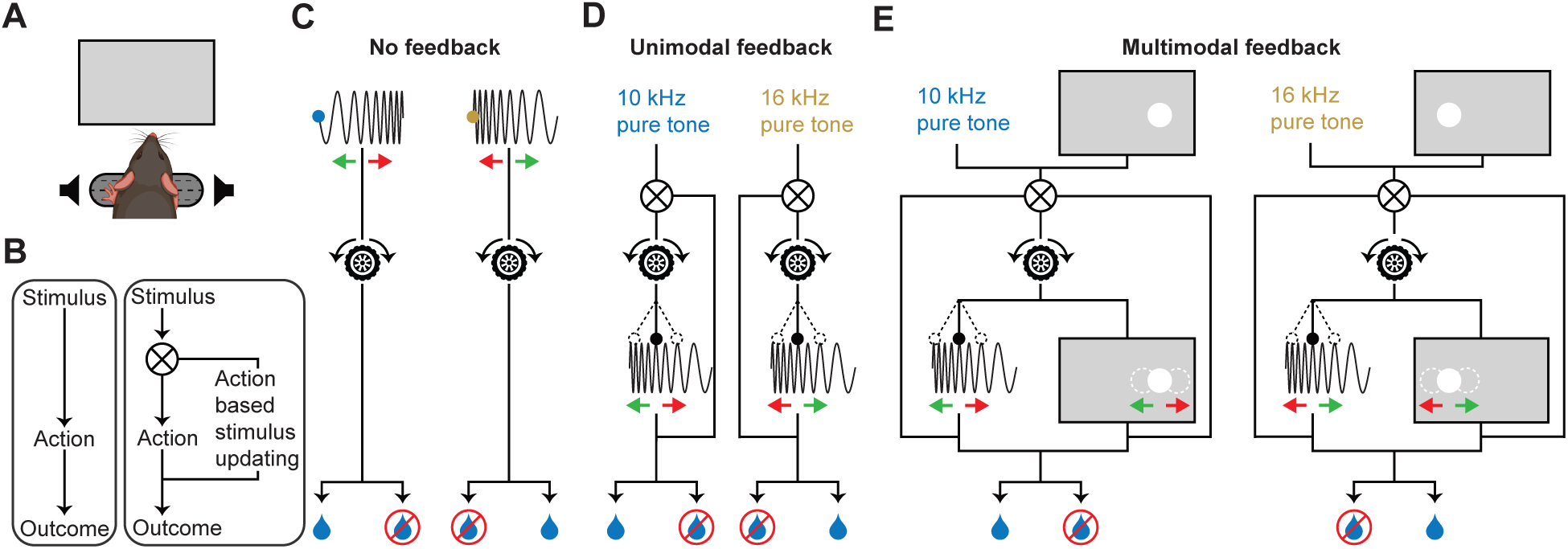
Experimental setup and closed-loop sensory feedback. **(A)** Schematic of a head-fixed mouse resting its front paws on a choice wheel. A computer monitor is placed in front of the mouse with a speaker on either side. **(B)** Overview of training without feedback (left) and with closed-loop feedback (right). **(C)** Schematic of training using no feedback. Auditory cues in the form of ascending (left) or descending (right) frequency modulated sweeps are presented without coupling to the movement of the wheel. Green arrows indicate the required direction of wheel movement to obtain water reward. **(D)** Schematic of training using unimodal auditory feedback. Leftward or rightward movements of the wheel result in the increase or decrease of the frequency of the auditory cue, respectively. Green arrows indicate the required direction of wheel movement to obtain water reward **(E)** Schematic of training involving multimodal auditory and visual feedback. Leftward or rightward movements of the wheel result in the increase or decrease of the frequency of the auditory cue as well as a horizontal displacement of the on-screen position of a visual cue to the left or right, respectively. Green arrows indicate the required direction of wheel movement to obtain water reward.

The major step in training sensory discrimination behaviors is to teach subjects to respond to task-relevant sensory stimuli with a volitional, targeted behavioral report (output). This step is time consuming in non-verbal animals, in which mice must link the required motor response to a preceding sensory cue by exploring all possible motor alternatives ^21–23^. In most training regimes ^24,25^, the sensory cue is non-interactive, i.e., motor responses do not elicit modifications of the sensory cue to indicate the relevance and importance of ongoing movement for the task at hand (**Fig. 1B**, left). If the consequences of motor responses are signaled only at the end a trial, a subject is unable to accumulate and use evidence from within the trial to guide movement choices and possibly make corrections. In contrast, providing intra-trial sensory feedback coupled to the motor response in a closed-loop fashion (**Fig. 1B**, right) can inform mice about the correctness of their ongoing movements. This allows accurate motor patterns to be confidently executed while erroneous patterns can be rectified in real-time.

In training regimes that use movement coupled sensory feedback ^13,26–29^, movement dynamically elicits changes to the sensory cue, i.e. a unique sensory cue is associated with each position of the response space. Hence, trained subjects have access to a library of sensorimotor associations from which they may invoke a desired motor pattern. All the above studies used movement coupled sensory feedback in a single sensory modality to indicate accuracy of ongoing movement. However, the contribution of feedback to learning and performance has not been systematically evaluated, as these studies employed feedback without directly manipulating its presence or characteristics.

In this study, we hypothesized that providing within-trial sensory feedback using a continuous response device would be beneficial during instrumental learning. A continuous response device affords the possibility to track decisions as they evolve within a trial and to use the current state of the response to provide sensory feedback; these features are unavailable in response devices which register decision reports as discrete events towards the end of a trial (e.g., lick left/right events). Continuous response devices have their limitations; they might be challenging to use because they require a novel motor response that is not in the native behavioral repertoire of the subject (e.g., a mouse moving a wheel to the left/right using its forelimbs to report choices). Such a device would require subjects to exploratively discover the end points of a behavioral report i.e. determine the implicit thresholds of a behavioral report by trial and error. Thus, it would be desirable to implement training procedures such as providing multimodal feedback to maximize the benefit of using continuous response devices.

Here, we trained three cohorts of mice on a sensory discrimination task with different feedback regimes: animals received either unimodal sensory (auditory) feedback, multimodal sensory (visual and auditory) feedback, or no feedback at all. We found that animals that received auditory feedback were quicker to acquire instrumental associations and were more accurate in trials with choice modifications compared to no-feedback controls. Further, we found that intratrial cumulative wheel movements of feedback animals, but not animals without feedback, underwent refinement as instrumental learning progressed. Our study thus shows that closed-loop sensory feedback can be used as a potent training aid to enable mice to acquire instrumental associations in a reproducible and time efficient manner.

## RESULTS

### Closed-loop sensory feedback training regime

We used a modified version of the choice wheel ^13^ to train head fixed mice to perform targeted wheel movements in response to sensory cues. We trained a total of 38 mice in 761 sessions (46,397 trials) on a two-alternative forced-choice task, in which animals indicated their responses to a sensory cue (auditory or audiovisual) by turning the choice wheel with their front paws either to the left or to the right.

The first cohort (n = 18) received no feedback, i.e. wheel movements were not coupled to changes of the auditory cue (**Fig. 1C**). The second cohort (n = 16) received unimodal auditory feedback, i.e. wheel movements resulted in changes of the auditory cue (**Fig. 1D**). The third cohort (n = 4) received multimodal auditory and visual feedback, i.e. wheel movements were coupled to changes of both auditory and visual cues (**Fig. 1E**). In addition to the auditory feedback, which was the same as for the second cohort, animals were presented a visual cue whose on-screen position was coupled to the position of the wheel.

### Animals receiving feedback reach the expert stage faster and more reliably

Mice in all training regimes learned to move the wheel to a predetermined threshold to report their choice after cue onset (see exemplar trials in **Fig. 2A**). While only 9 out of 18 (50 %) animals trained with no feedback achieved a task performance greater than 70 % (categorized as ‘task acquired’), 10 out of 16 (62 %) animals trained with unimodal feedback and 4 out of 4 (100 %) animals trained with multimodal feedback acquired the task.

**Figure 2:**
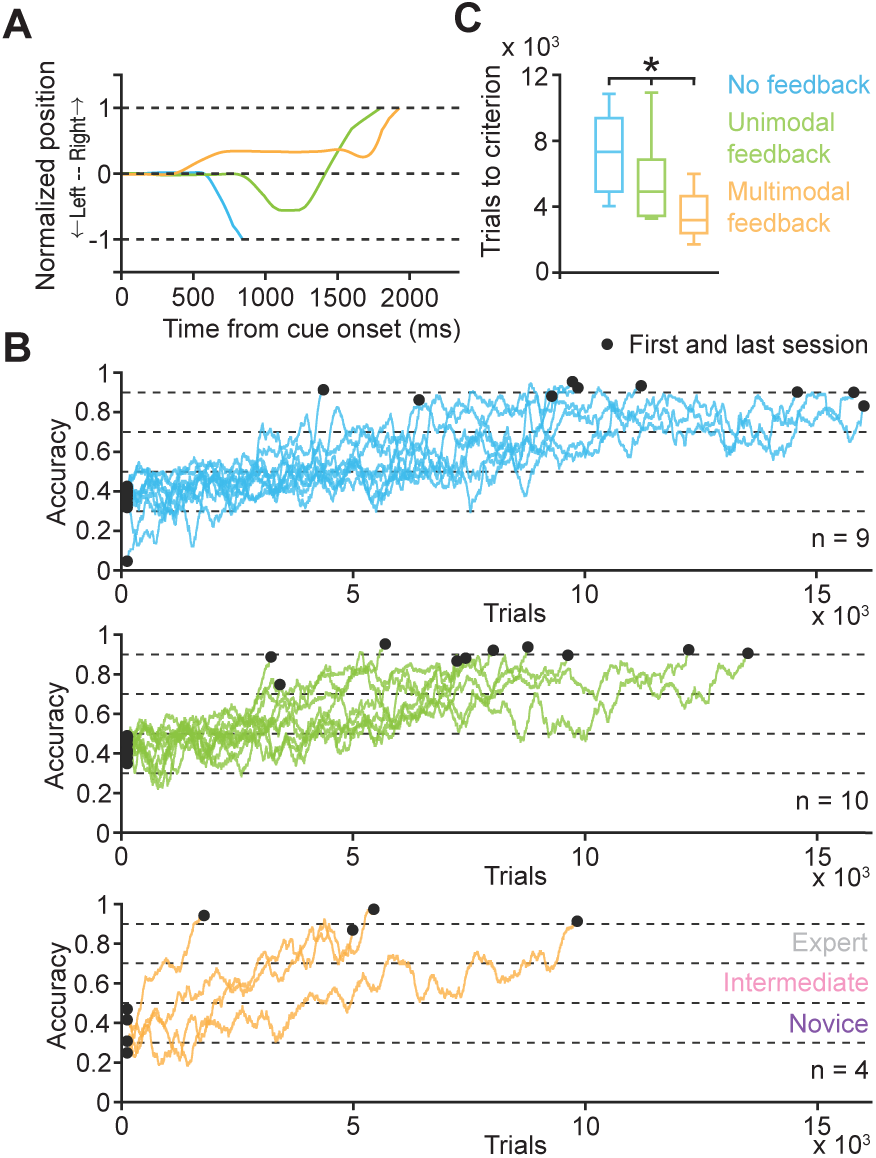
Animals receiving feedback reach the expert stage faster. **(A)** Representative movement traces from each training cohort. Dotted lines indicate threshold for trial termination. **(B)** Learning curves of individual mice that successfully acquired the task in the no feedback (n = 9), unimodal feedback (n = 10) and multimodal feedback (n = 4) cohorts. Task performance is classified into novice (30-50 %), intermediate (50-70 %) and expert (70-90 %) stages. Black dots indicate the first and last training session. **(C)** Median number of trials across animals in each cohort required to reach the expert stage. Kruskal Wallis test, p = 0.0245.

We categorized the animals’ task performance over the course of training into novice (30-50 %), intermediate (50-70 %) or expert stages (70-90 %) (**Fig. 2B**). Animals in the three training regimes significantly differed in the number of trials required to reach the expert stage; animals that received feedback required fewer trials to reach the expert stage compared to animals that did not receive feedback (**Fig. 2C**) (median number of trials, 7329 no feedback vs. 4918 unimodal feedback vs. 3186 multimodal feedback. Kruskal Wallis test, p = 0.0245). In other words, unimodal feedback and multimodal feedback cut down training duration to 67 % and 43 %, respectively, compared to no feedback. Notably, the fastest learner in the multimodal feedback cohort reached the expert stage faster than the fastest learner in either of the other two cohorts. Even the slowest learner in the multimodal feedback cohort still reached the expert stage faster than the slowest learners in the no feedback and unimodal feedback cohorts (see **Fig. 2B**).

Next, to evaluate the level of task engagement across training in the three training regimes, we computed the fraction of total trial duration in which the wheel was not moved. We found that the no-movement fraction significantly decreased across task acquisition in all three cohorts (Kruskal Wallis test: p = 6.41×10^−8^ no feedback; p = 2.05×10^−7^ unimodal feedback; p = 1.17×10^−11^ multimodal feedback) (**Fig. 3**). Taken together, the results show that although mice in all cohorts become increasingly task engaged across task acquisition, providing closed-loop sensory feedback is beneficial for accelerated and reliable task acquisition.

**Figure 3:**
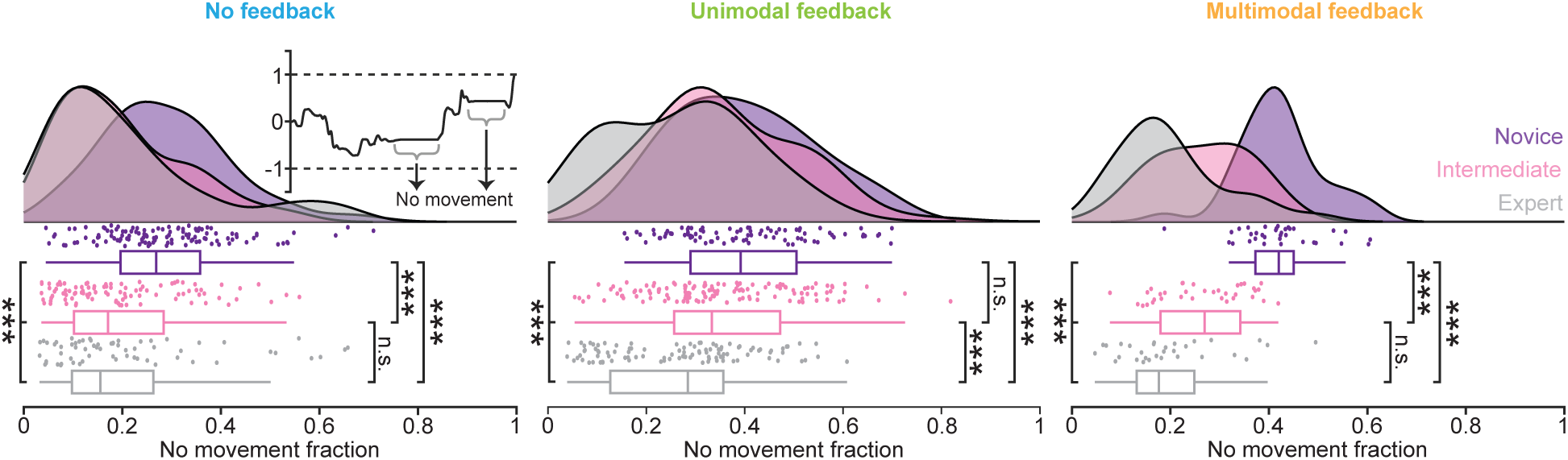
Progress of instrumental learning is accompanied by increased task engagement. Raincloud plots showing the probability distribution, individual session values (dots) and box plots for no movement fraction across training stages in the three cohorts. Box plots show median (central line), interquartile range (box boundaries), and data range (whiskers). Novice, intermediate and expert training stages are color coded. Left, no feedback (Kruskal-Wallis test, p = 6.41×10⁻⁸). Middle, unimodal feedback (Kruskal-Wallis test, p = 2.05×10⁻⁷). Right, multimodal feedback (Kruskal-Wallis test, p = 1.17×10⁻¹¹). Inset shows an exemplar trial with movement and no-movement epochs. Pairwise comparisons are conducted using the post-hoc Dunn test. *, p < 0.05, **, p < 0.01, ***, p < 0.001.

### Feedback increases the occurrence and accuracy of choice-modified trials

To elucidate the behavioral signatures that accompany accelerated task acquisition, we investigated single-trial wheel movement trajectories in detail. Since the number of trials required to reach criterion accuracy were significantly different amongst the three cohorts, we hypothesized that wheel movements in each cohort might be qualitatively different. Analysis of wheel movements after cue presentation in all cohorts revealed two types of correct trials: trials involving directed movements to the correct threshold and trials involving initial erroneous movements that were subsequently modified to produce a movement directed at the correct threshold (i.e. trials with ‘flips’ or choice modifications) (**Fig. 4A, B**). We defined a flip as a change in the direction of wheel movement after the wheel had been moved a distance of more than half the current session threshold (**Fig. 4B**, middle, right). In all training stages, animals without feedback displayed the smallest proportion of correct trials with flips; animals with feedback displayed a significantly greater proportion of correct trials with flips than animals without feedback (expert stage: 17 % multimodal feedback, 11 % unimodal feedback, 9 % no feedback; Kruskal Wallis test, p = 1.04×10^−6^) (**Fig. 4C**). This result suggests that providing feedback promotes the execution of trials where corrective movements are involved.

**Figure 4:**
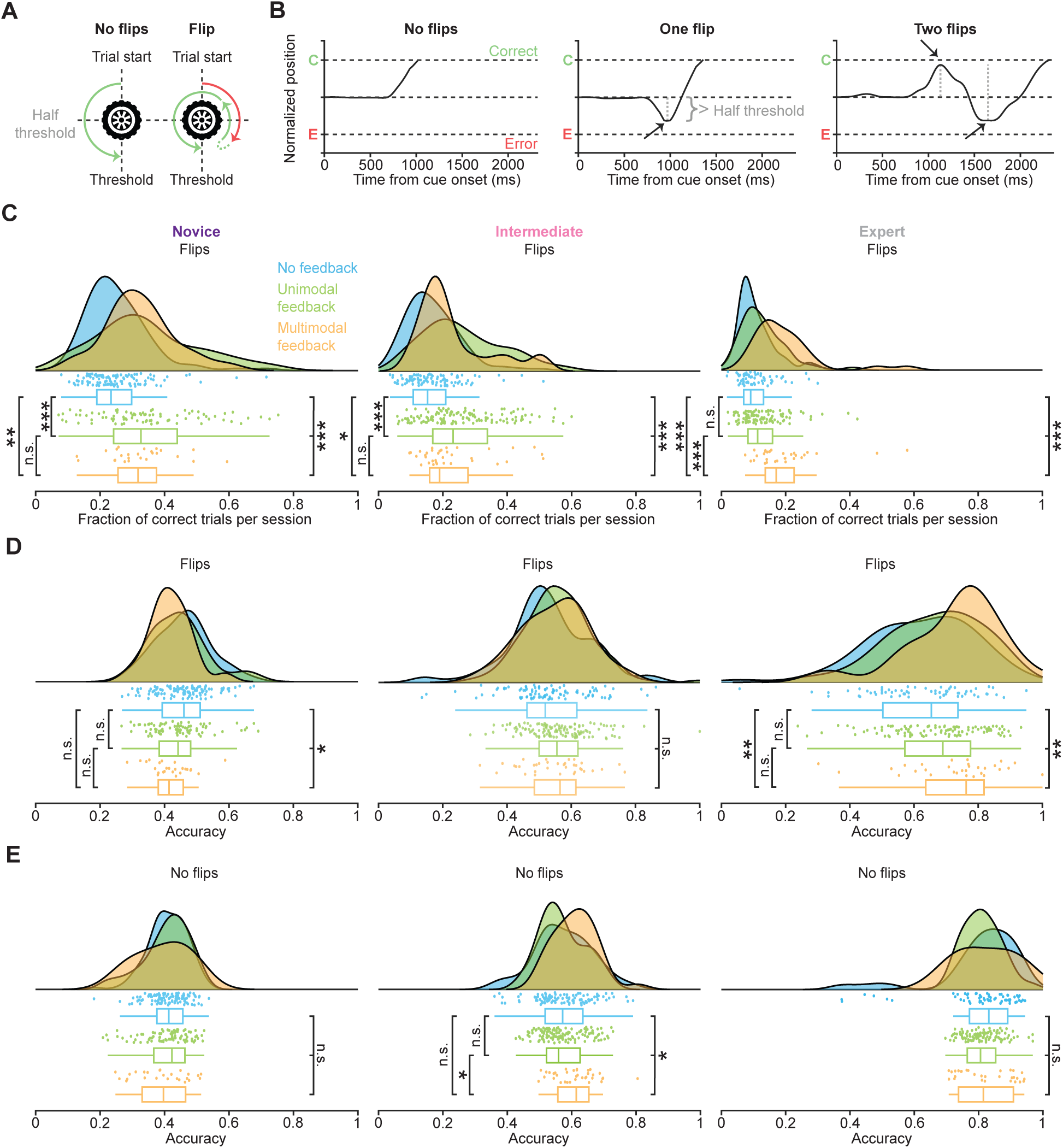
Feedback increases the occurrence and accuracy of choice-modified trials. **(A)** Schematic of trials without and with choice modifications (flips). Left, a trial where the wheel is moved to the threshold with a single, directed movement. Right, the wheel is initially moved towards the erroneous side, and the movement is subsequently rectified by moving the wheel towards the correct side. Green and red arrows indicate correct and incorrect movements, respectively. **(B)** Representative movement trajectories in trials without and with choice modifications. Left, Exemplar trial in which the wheel is moved to the threshold with a single, directed movement, i.e. without a flip. Middle, a trial with a single flip. Right, a trial with two flips. Vertical dotted lines highlight the extent of movement before a flip is initiated. **(C)** Distribution of the fraction of correct flip trials for novice, intermediate and expert training stages. Raincloud plots showing the probability distribution, individual session values (dots) and box plots; box plots show median (central line), interquartile range (box boundaries), and data range (whiskers). Left, novice stage (Kruskal-Wallis test, p = 2.92×10⁻⁶). Middle, intermediate stage (Kruskal-Wallis test, p = 1.47×10⁻⁸). Right, expert stage (Kruskal-Wallis test, p = 1.04×10⁻⁶). Pairwise comparisons are conducted using the post-hoc Dunn test. **(D)** Same layout as (C) plotting the distribution of the accuracy of flip trials in a session. Left, novice stage (Kruskal-Wallis test, p = 4.42×10⁻²). Middle, intermediate stage (Kruskal-Wallis test, p = 1.84×10⁻¹). Right, expert stage (Kruskal-Wallis test, p = 5.36×10⁻³). **(E)** Same layout as (C) plotting the distribution of the accuracy of trials involving no flips in a session. Left, novice stage (Kruskal-Wallis test, p = 7.31×10⁻¹). Middle, intermediate stage (Kruskal-Wallis test, p = 1.62×10⁻²). Right, expert stage (Kruskal-Wallis test, p = 1.99×10⁻¹). *, p < 0.05, **, p < 0.01, ***, p < 0.001.

Next, we asked whether the accuracy of flip trials evolved differently across training for the feedback and no feedback cohorts. In the expert stage, feedback animals displayed significantly higher accuracy in trials with flips compared to animals without feedback (76 % multimodal feedback, 68 % unimodal feedback, 65 % no feedback; Kruskal Wallis test, p = 5.36×10^−3^) (**Fig. 4D**). This effect was not apparent in the novice training stage, suggesting that feedback animals gradually develop clarified instrumental associations that aid in successful execution of flip trials. Remarkably, we found that accuracy in trials without flips were not significantly different across the training stages between the feedback and non-feedback cohorts (expert stage: 81 % multimodal feedback, 80 % unimodal feedback, 83 % no feedback; Kruskal Wallis test, p = 0.199) (**Fig. 4E**). Taken together, these findings argue that feedback selectively enables mice to successfully conduct wheel movements that serve to correct initial erroneous choices.

### Feedback refines wheel movements across training

Next, we investigated whether progress in task acquisition was accompanied by a refinement in the cumulative wheel movements of individual trials as a measure of motor precision. We defined pathlength as the distance traversed by the wheel during a trial regardless of movement direction; for this measure, epochs of the trial where the wheel was not moved, were excluded. While pathlength in flips trials across training stages remained unchanged for the no-feedback cohort (Kruskal Wallis test, p = 0.2602) (**Fig. 5A**, left), it significantly decreased across learning for the feedback cohorts (Kruskal Wallis test, p = 4.07×10^−18^ unimodal feedback; p = 8.02×10^−6^ multimodal feedback) (**Fig. 5A**, middle and right). Further, we found that pathlength in no-flip trials significantly decreased in all cohorts and, they were shorter, suggestive of higher confidence in this trial type (Kruskal Wallis test, p = 5.83×10^−11^ no feedback; p = 3.88×10^−18^ unimodal feedback; p = 4.09×10^−7^ multimodal feedback) (**Fig. 5B**). These results indicate that a refinement of motor output accompanied task acquisition selectively in flip trials in the feedback cohorts; unchanging pathlengths in the no-feedback cohort indicate that mice moved the wheel with lesser clarity about task contingencies, a feature which was also reflected in the lower accuracy of flip trials (**Fig. 4D**).

**Figure 5:**
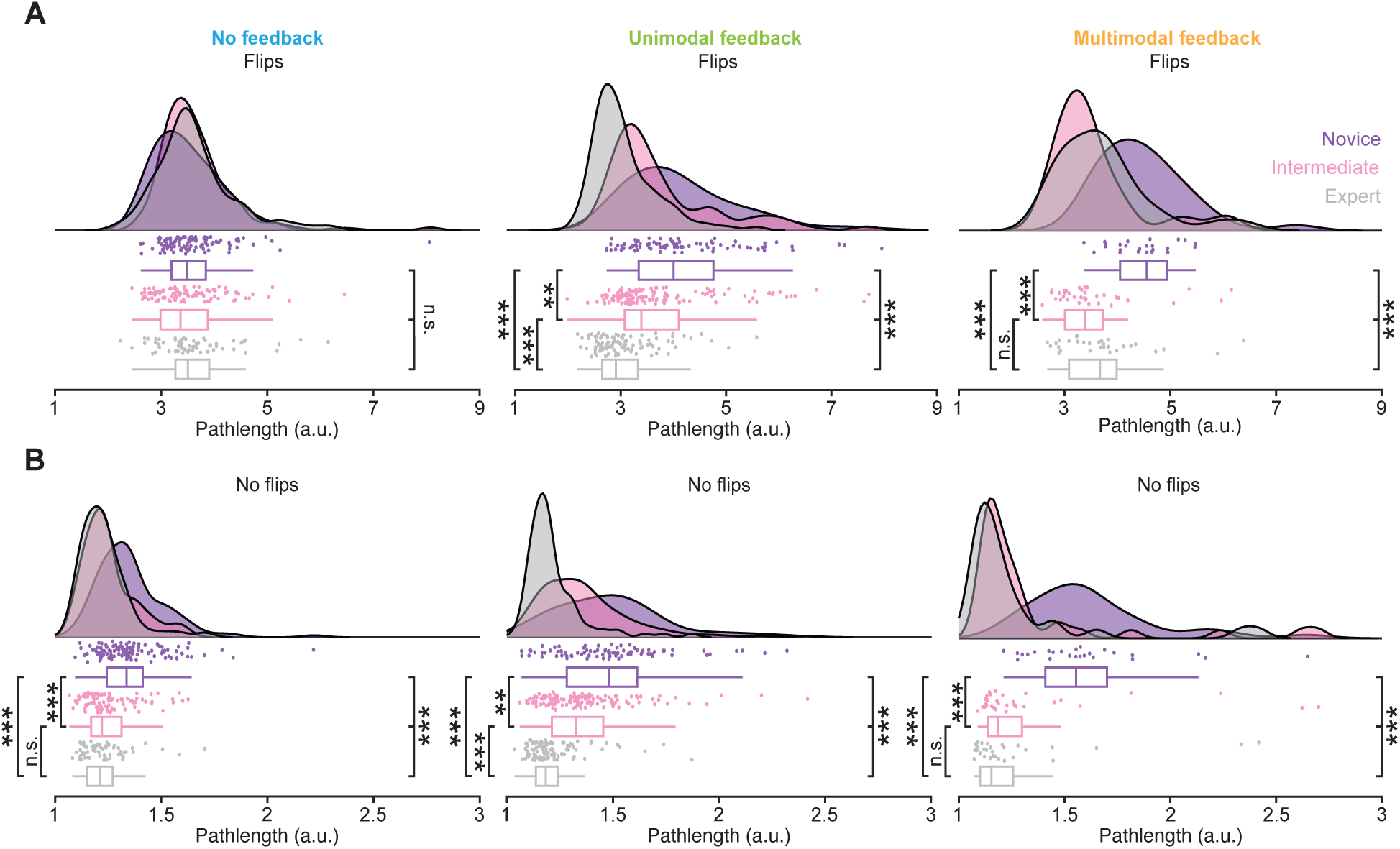
Feedback refines wheel movements across training. **(A)** Distribution of pathlength in flips trials. Raincloud plots showing the probability distribution, individual session values (dots) and box plots; box plots show median (central line), interquartile range (box boundaries), and data range (whiskers). Left, no feedback (Kruskal-Wallis test, p = 2.60×10⁻¹). Middle, unimodal feedback (Kruskal-Wallis test, p = 4.07×10⁻¹⁸). Right, multimodal feedback (Kruskal-Wallis test, p = 8.02×10⁻⁶). Pairwise comparisons are conducted using the post-hoc Dunn test. **(B)** Same layout as (A) showing the pathlength in trials without flips. Left, no feedback (Kruskal-Wallis test, p = 5.83×10⁻¹¹). Middle, unimodal feedback (Kruskal-Wallis test, p = 3.88×10⁻¹⁸). Right, multimodal feedback (Kruskal-Wallis test, p = 4.09×10⁻⁷). Note the different scaling in (B). *, p < 0.05, **, p < 0.01, ***, p < 0.001.

### Animals unsuccessful in task acquisition exhibit erroneous and shorter exploratory wheel movements

Finally, we sought to determine whether mice that were unsuccessful in acquiring task contingencies (n = 9, no feedback; n = 6, unimodal feedback) differed in the behavioral signatures investigated above in the formative stages of instrumental associations, i.e. in the novice training stage. In comparison to successful mice, unsuccessful mice in the no-feedback and unimodal feedback cohorts displayed a significantly lower proportion of correct trials involving choice modifications (Rank-sum test, p = 6.24×10^−4^ no feedback; p = 9.91×10^−4^ unimodal feedback) (**Fig. 6A, B**). Further, pathlengths in flip trials were shorter for unsuccessful mice (Rank-sum test, p = 0.0152 no feedback; p = 0.0179 unimodal feedback) (**Fig. 6A, B**). Health scores (**Fig. S1**), weight and water intake (**Fig. S2**) of animals that did not acquire the task were similar to those of animals that successfully acquired the task. Together, these results suggest that the lower fraction of correct flip trials driven by a lack of exploratory wheel movements in the novice stage could underlie the failure to acquire task contingencies in unsuccessful mice.

**Figure 6:**
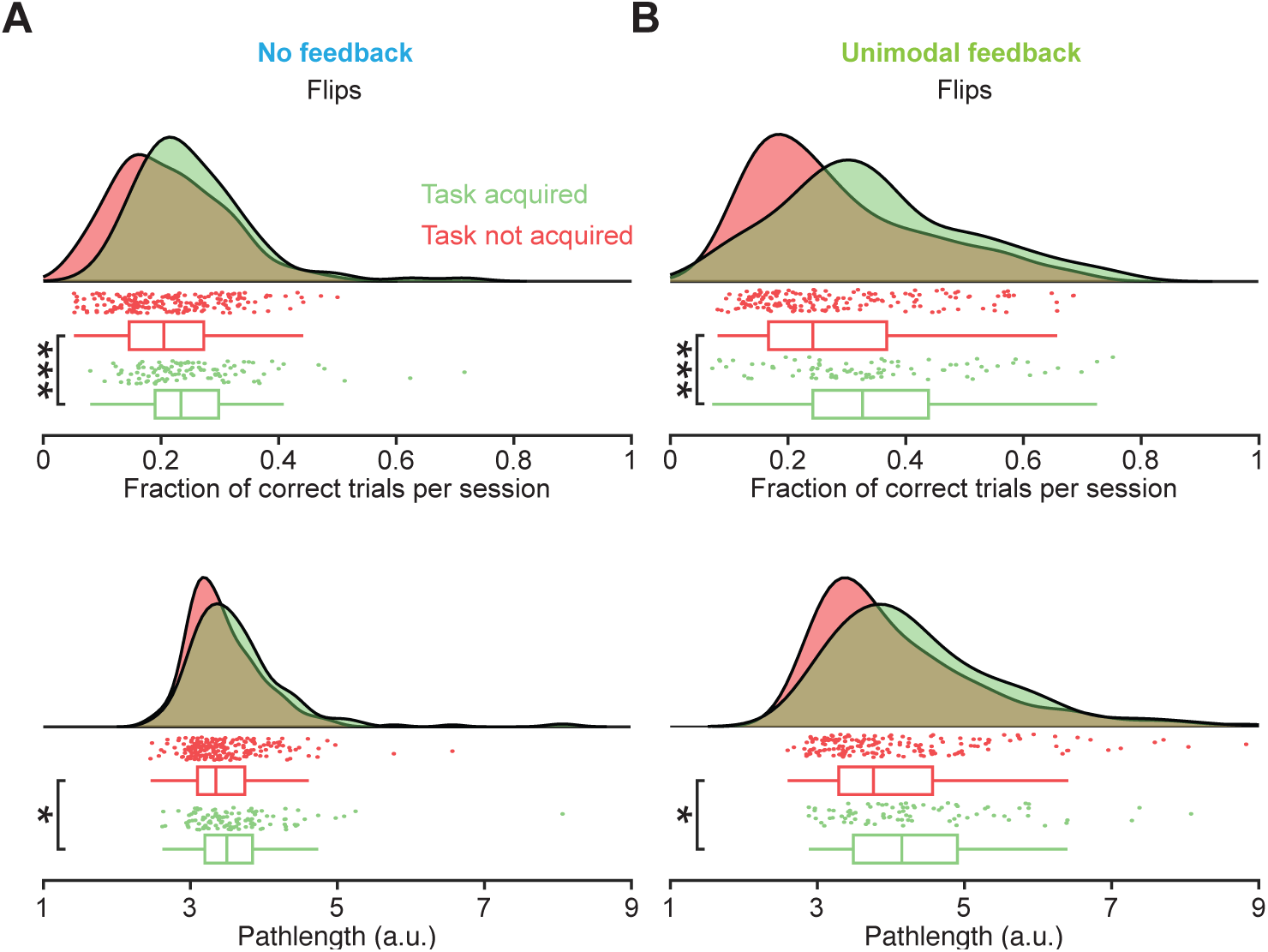
Animals unsuccessful in task acquisition exhibit erroneous and shorter exploratory wheel movements. **(A)** Fraction of correct flip trials and pathlengths in flips trials comparing ‘task acquired’ and ‘task not acquired’ animals for the no feedback cohort. Raincloud plots showing the probability distribution, individual session values (dots) and box plots; box plots show median (central line), interquartile range (box boundaries), and data range (whiskers). Top, fraction of correct flips trials per session. Wilcoxon rank-sum test, p = 6.24×10^−4^. Bottom, median pathlength in flips trials per session. Wilcoxon rank-sum test, p = 1.52×10^−2^ **(B)** Same layout as (A) for the unimodal feedback cohort. Top, fraction of correct flips trials per session. p = 9.91×10^−4^. Bottom, pathlength in flips trials per session. Wilcoxon rank-sum test, p = 1.79×10^−2^. *, p < 0.05, ***, p < 0.001

## DISCUSSION

Our work shows that animals provided with closed-loop sensory feedback acquire instrumental associations faster and with clearer task contingencies. The mechanism underlying this advantage was revealed by analyzing trials with choice modifications: feedback animals showed progressive increases in the proportion and accuracy of choice modified trials, along with refined wheel movements. This suggests feedback animals better grasped task contingencies by leveraging real-time sensory information to re-evaluate and correct erroneous movements within trials, a possibility unavailable to non-feedback animals that could only learn from end-of trial outcomes.

The choice wheel, an increasingly used response device to study decision making in mice ^30^, facilitates continuous tracking of the motor manifestations of evolving decision states that can be used to provide feedback about the accuracy of ongoing movement within a trial. Studies investigating the serial order of decision formation and movement execution have indicated that an evolving decision is expressed in behavior even before it is finalized ^31,32^. Further, modeling studies have indicated that the brain continues to process incoming sensory information even after an initial decision has been made ^33^. Thus, using a continuous response device such as the choice wheel allows experimenters to describe ongoing motor responses as representatives of instantaneous decision states shaped by hitherto accumulated evidence; these decision states can then be influenced by supplying feedback to ensure successful outcomes. In comparison, using non-continuous, discrete decision reports such as lick events, nose pokes and lever presses limit experimenters to register only the endpoint of a decision, thus preempting intra-trial feedback. The current study maximizes the usage of the choice wheel by not only using wheel position to describe evolving decision states but also by using it to provide feedback in multiple sensory modalities. The implementation of such standardized training paradigms in experiments involving the choice wheel would significantly contribute toward increasing the efficiency and reproducibility of instrumental training.

The training advantage conferred by closed-loop sensory feedback can be better comprehended when voluntary movements are conceptualized through the forward and inverse models of motor control ^34^. Forward models compute the sensory consequences of intentional movements by comparing the actual position of the body with its predicted position. Forward models thus serve to inform an organism of deviations from predicted bodily sensory percepts after executing planned movements. On the other hand, inverse models generate motor commands based on intentions to move the body to a desired location. Together, the inverse and forward models provide a conceptual framework for the generation and refinement of voluntary movements. In the context of instrumental learning with sensory feedback, we speculate that the refinement of the inverse model is driven by minimizing the output of the forward model i.e., by reducing the mismatch between expected and actual sensory percepts resulting from intentional movements. As training progresses, a subject can execute movements with increasingly predictable sensory consequences due to the formation of clarified sensorimotor associations and use it to ensure successful outcomes. Concretely stated, an expert mouse successfully implements choice modifications because it would be able to accurately predict changes to the sensory cue. A study ^35^ examined whether humans use sensory feedback (forward model) or cognitive strategy (inverse model) when making visually guided movements with directional perturbation. Subjects were instructed to counteract perturbations using a fixed offset angle. However, as they practiced, they increasingly relied on continuous visual feedback rather than the fixed cognitive strategy. This suggests that task proficient subjects executed movements in alignment with what they saw rather than employ an explicit cognitive strategy. These findings are in alignment with our results in showing that the increased predictability of the sensory consequences of movement (i.e. minimized sensory deviations) might provide an accurate estimate of task contingencies.

The neural dynamics accompanying multisensory task input and its behavioral significance have been highlighted by recent electrophysiological studies in different species. In mice, frontal sub-regions like the secondary motor cortex additively combine incoming auditory and visual task signals and this process appears to evolve with task acquisition ^36^. For our study, this finding has the implication that task acquisition in animals with multimodal feedback is likely quicker due to a richer representation of the sensory environment in an executive brain region, which enables better decision making. It has been demonstrated that providing artificial sensory feedback by means of intracortical microstimulation of the primary somatosensory cortex can be used by non-human primates ^37^ and humans ^38^ to generate accurate motor sequences. Further, congruent coupling of motor output and visual input is shown to be necessary for normal functional development of layer 2/3 neurons in the primary visual cortex of mice ^39^. These studies indicate that specific neural circuits are in operation when sensory feedback is used to guide movements. Longitudinal monitoring of activity in these circuits can throw light on the neural processes accompanying feedback driven decision making and subsequent movement execution.

The behavioral trends observed in the multimodal feedback cohort, though based on a limited sample size (n = 4 animals), are consistent with the beneficial effects of multimodal sensory stimulation previously reported in mice using comparably small samples. A similarly powered study (n = 7 animals) showed that mice in a detection task exhibited higher detection accuracy for multimodal compared to unimodal stimuli, with response rates improving by approximately 25 % in the multimodal condition and reaction times being significantly shorter than with unimodal cues alone^40^. Similarly, another study (n = 9 animals) demonstrated in an operant paradigm that paired multimodal stimuli yielded higher response accuracy than either modality alone across all tested stimulus durations, with the greatest gains observed at shorter stimulus presentations^41^. Importantly, both studies included unimodal visual controls and demonstrated that audiovisual advantages exceeded those of visual stimulation alone, suggesting that the benefits of multimodal feedback observed in the present study reflect genuine cross-modal enhancement rather than a contribution of visual feedback alone. Further, scoping reviews and meta-analyses of human studies confirm that multimodal stimuli confer processing advantages, i.e., they are consistently perceived faster and retained longer than unimodal cues^42,43^.

In conclusion, the current study demonstrates that closed-loop sensory feedback leads to quicker instrumental acquisition by clarifying task contingencies. Incorporating sensory feedback during instrumental acquisition can lead to quicker and more reliable training of experimental subjects and thus improve reproducibility of neurobehavioral experiments.

## SUPPLEMENTARY FIGURES

**Figure S1:**
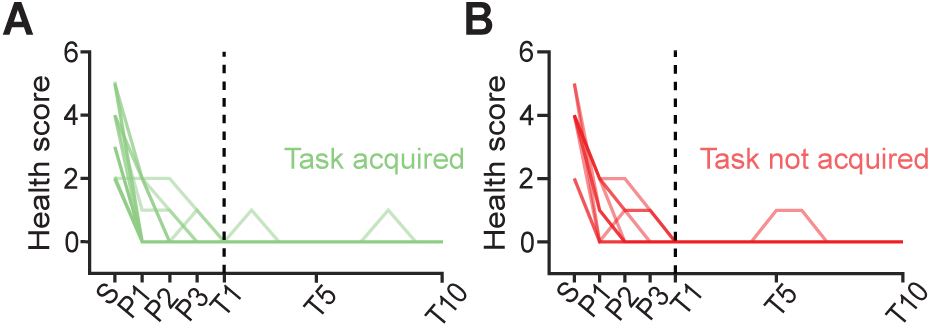
Health scores of animals that acquired or did not acquire the task. **(A)** Scores of animals that successfully acquired the task. Health scores incorporated assessments of activity, posture, grooming and signs of dehydration. Lower scores indicate better health. **(B)** Same layout as (A) for animals that did not acquire the task. S = Day of surgery, P1 = Post-operative day 1, P2 = Post-operative day 2, P3 = Post-operative day 3, T1 = Training day 1, T5 = Training day 5, T10 = Training day 10.

**Figure S2:**
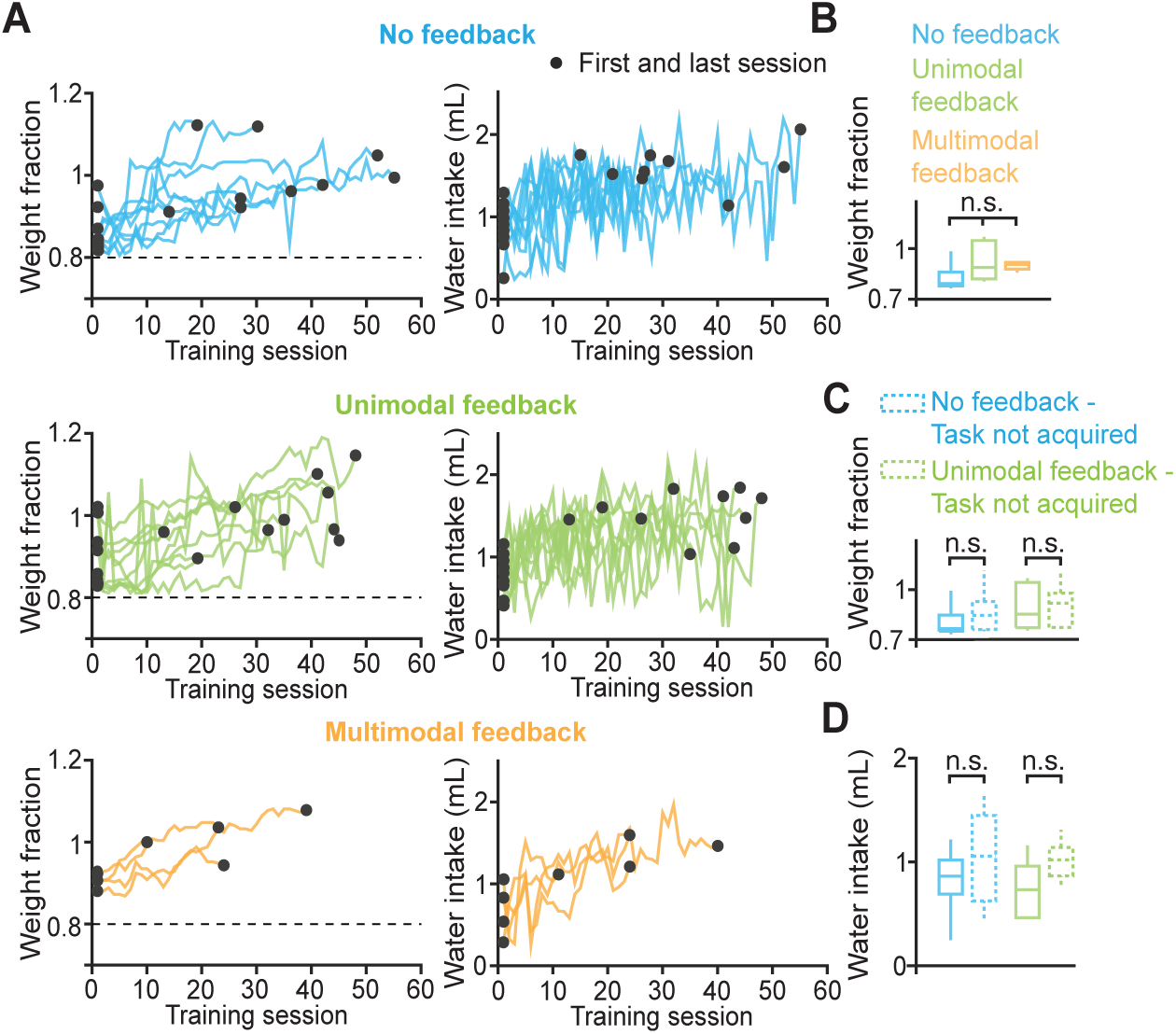
Weights and water intake among cohorts. **(A)** Left column, individual animal weights (fraction of pre-surgery weight) in the no feedback (top), unimodal feedback (middle) and multimodal feedback (bottom) cohorts. Right column, individual animal water intake in the no feedback (top), unimodal feedback (middle) and multimodal feedback (bottom) cohorts. **(B)** Median fraction of pre-surgery weights on the first day of training in the no feedback, unimodal feedback and multimodal feedback cohorts. Kruskal-Wallis test, n.s., not significant. **(C)** Median fraction of pre-surgery weight on the first day of training in the task acquired (solid lines) and task not acquired (dotted lines) groups in the no feedback and unimodal feedback cohorts. Wilcoxon rank-sum test, n.s., not significant **(D)** Median water intake on the first day of training in the task acquired (solid lines) and task not acquired (dotted lines) groups in the no feedback and unimodal feedback cohorts. Wilcoxon rank-sum test, n.s., not significant.

## MATERIALS AND METHODS

### Animals

A total of 38 male C57BL/6J mice sourced from Charles River (France) were used for behavioral training. All animal procedures were authorized by the local government (Regierung von Oberbayern, license number ROB-55.2-2532.Vet_02-17-119). Animal health was examined and scored every day. Wild-type male mice (C57BL/6J, Charles River) were used for all experiments. Mice were 8–10 weeks old at the beginning of the experiments and were housed in single cages on a reversed 12-h light/12-h dark cycle. Ambient humidity and temperature were set to 50 % and 24°C, respectively. Mice had ad libitum access to food and water except during behavioral experiments.

### Behavioral setup

Behavioral training was conducted in sound-attenuated operant chambers (Med Associates, USA) where components necessary for training were mounted using an optical breadboard (Thorlabs, USA). Mice were head-fixed at an elevation of 10 cm and placed in a plastic tube to allow free body movements. A 10-inch LCD screen (25.4 cm diagonal; Faytech FT10TMB (resolution 1920×1080), Germany) was mounted 15 cm in front of the animals to signal trial start and deliver visual stimuli. Electrostatic speakers capable of ultrasonic production (ES1; Tucker Davis Technologies, USA) were positioned 10 cm to the left and right of the animals’ ears to deliver auditory cues. Water reward was delivered using a spout placed within reach of the tongue, connected to a 5 ml disposable syringe driven by a TTL-controlled pump (NE-500, New Era Pump Systems, USA). Choice wheel movements were registered using a data acquisition device (PCIe-6323, National Instruments, USA) with a breakout panel (BNC-2090A, National Instruments, USA). The behavioral task was administered using custom MATLAB code and MonkeyLogic 2 ^44^, a NIMH toolbox running in MATLAB 2018a (Mathworks, USA).

### Trial Structure

A trial began with the monitor screen in front of the head-fixed mice turning gray. Mice were required to hold the choice wheel still for 1 s. After successfully holding the wheel still for 1 s, mice were presented with either no feedback or feedback sensory cues. After cue onset, wheel movements were registered for a maximum period of 60 s after which the trial would be terminated. Wheel movements during this period that crossed a predetermined threshold were categorized as correct or incorrect depending on the presented sensory cue. If no threshold crossings were detected, the trial would be marked as a missing trial. Correct trials ended with a water reward while error and missing trials ended with an omission of water reward. Trial completion was followed by an inter-trial interval (ITI) of 5 s.

### Sensory Feedback

#### Auditory feedback

Mice were presented a pure tone whose frequency was coupled to the direction of wheel movement. Leftward or rightward movements resulted in an increase or decrease of the frequency of the presented pure tone, respectively. That is, 1 degree of wheel movement resulted in 88.8 Hz increase or decrease in the frequency of the presented pure tone. Go-left trials commenced with the presentation of a 10 kHz pure tone and terminated when the frequency reached 14 kHz or 6 kHz for a correct or incorrect trial respectively. Go-right commenced with the presentation of a 16 kHz pure tone and terminated when the frequency reached 12 kHz or 20 kHz for a correct or incorrect trial respectively. The choice of the starting frequencies of the pure tones in the go-left and go-right trial types were motivated by the starting frequencies of the FM sweeps in the go-left and go-right trials in the no feedback regime.

To implement closed loop coupling of steering wheel movement and auditory sensory cues, movement data from the wheel (i.e. voltage values mapping to wheel rotations) was fed into an auditory processor (Tucker-Davis Technologies RX8). On the auditory processor a custom written program generated a sine waveform whose frequency was linearly mapped to wheel-derived voltage values. Mapping of wheel rotation and sine frequencies was done as follows.

The range of possible voltage values from the wheel rotation was determined by the movement thresholds. The range of voltage values were then mapped to a range of sine frequencies i.e. each position of the wheel was associated with a unique sine frequency. During a trial, wheel movements which modulated the sine frequencies and produced by the processor were converted to sounds by the speakers.

#### Visual feedback

To implement closed loop coupling of steering wheel movement and visual feedback cues, movement data from the wheel was fed into MonkeyLogic, which controlled the on screen rendering of a white filled circle (RGB: 1, 1, 1) with a radius of 10 degrees. Similar to the auditory feedback, the possible voltage values from wheel rotation was determined by the movement thresholds. Extremes of the voltages (representing the thresholds) were mapped to the extremes in the horizontal position of the visual cue. The visual extremes were set as the left border and the center of the screen, and the right border and the center of the screen, for go-right and go-left trials, respectively. Thus, only one horizontal half of the screen was used per trial. Horizontal positions dependent on the wheel rotation were linearly interpolated between the extremes. Concretely, a trial started with the wheel in the neutral position and the visual cue in the middle of the current trial’s horizontal range (0.25 * screen width for go-right, 0.75 * screen width for go-left). A change of the wheel rotation linearly changed the cue’s position. Accordingly, rotating the wheel to the correct side horizontally moved the stimulus towards the screen center (0.5 * screen width), while rotating to the wrong side moved it towards the screen borders (0 * screen width and 1 * screen width for go-right and go-left, respectively).

#### No feedback

Go-left trials, in which supra-threshold wheel movements to the left side were rewarded, commenced with the presentation of a 1 s frequency modulated (FM) sweep ranging from 10 kHz – 14 kHz (78 dB SPL; 4 Hz/s). Go-right trials, where supra-threshold wheel movements to the right side were rewarded, commenced with the presentation of a 1 s FM sweep ranging from 16 kHz – 12 kHz (78 dB SPL; 4 Hz/s). Each FM sweep was repeated till the end of a trial with a break of 0.5 s between repetitions. The FM sweeps were chosen such that the sweeps fell within the frequency range of comparable auditory sensitivity in mice ^45–47^; they have proven reliable for investigating auditory discrimination tasks as evidenced by robust and replicable behavioral performance in earlier studies ^48,49^

### Surgery and headplate Implantation

Mice were anesthetized using 1-2% isoflurane and installed in a stereotactic frame (Neurostar). An analgesic (Novaminsulfon 200 mg/kg body weight) was subcutaneously injected prior to surgery. Body temperature was maintained at around 37°C using a heating pad for the entire duration of the surgery. Hair above the skull was removed using a shaver. The skin above the skull was disinfected with 70% ethanol and a topical anesthetic was subcutaneously injected (2% Lidocaine). After excising the skin to expose the skull, the surface was thoroughly cleaned and subsequently roughened in preparation for headplate implantation. After correcting for tilt and scaling of the head using an automated software (Neurostar), the headplate (Luigs & Neumann, Germany) was implanted at the anterior aspect of the cranium using dental cement. After surgery, mice were allowed to recover for at least 3 days before the commencement of behavioral training.

### Behavioral training

After recovery of at least 3 days from surgery, mice were subjected to a water restriction schedule to increase motivation during behavioral training where water was the reward. Mice were given 1500 μl of water per day during the course of a training session (once per day). If a mouse consumed less than 1500 μl during a training session, the remaining volume was supplemented after the session. Animals were trained on all weekdays and on alternate weekends. On non-training days, mice received 1500 μl water in their home cage. Animals were scored daily and the body weight was measured to ensure that it does not drop below 80% of the weight prior to surgery.

After the commencement of water restriction, animals were handled in the palm of the hand for 2-3 days to acclimatize to the experimenter. On these days, animals were given 1500 μl of water via a pipette. Next, animals were head-fixed in the training setup and habituated to moving the choice wheel and drinking water from the spout. During these pre-training sessions, animals would receive a reward of 10 μl per trial for moving the choice wheel to a low threshold in either direction. Pre-training lasted for 2-3 sessions.

After pre-training, mice were trained to respond to either feedback or no feedback cues. Each trial commenced with the presentation of a gray screen (RGB: 0.3, 0.3, 0.3) on the monitor in front of the animals; the gray screen was used to indicate trial start. The gray screen was kept on for the duration of the trial. After the presentation of the gray screen, animals were expected to not move the choice wheel for at least 1000 ms before the presentation of feedback or no feedback cues. A trial was terminated upon registering a thresholded response to the left or right side. Maximum trial duration was 30 s. Animals started training with a threshold of 20-25 degrees and progressed to 45 degrees as accuracy increased.

### Data analysis

All analyses were conducted using custom written software in MATLAB.

#### Detection of flips

Wheel movement data for each trial was normalized such that data in a given trial had a value of 0 at cue onset and reached −1 or 1 when the threshold was reached on the left/right side, respectively. Normalized wheel movement data from each trial was then input to the MATLAB function ‘findpeaks’ to detect local maxima, which reflect choice modifications. The parameter ‘MinPeakProminence’ which specifies the minimum value of the local maxima to be reached before it is counted as a peak was set at 0.5, i.e. a movement excursion had to be higher than 0.5 times the trial threshold to be counted as a choice modification/flip. After detection of flips in one direction, data from the same trial was sign inverted and fed into ‘findpeaks’ to detect flips in the opposite direction. Lastly, the local maxima count from the two iterations were summed to arrive at the total flips in that trial.

#### No movement fraction

To determine the periods in a trial during which the wheel had been held static, we took the first derivative of the wheel position and computed the instantaneous velocity for the entire trial. Next, we quantified the periods where the velocity was zero, i.e. the periods where the wheel was moved less than 0.02 degrees/s. We then divided the periods of zero velocity by the total trial duration which gave us the fraction of trial time where the where there was no wheel movement.

### Statistical analyses

Data were analyzed using a Kruskal Wallis test followed by a post-hoc Dunn test or a Wilcoxson rank-sum test. The following significance levels were used in the manuscript: *, p < 0.05, **, p < 0.01, ***, p < 0.001.

## DATA AND CODE AVAILIBILITY

All data and code from this paper will be shared by the lead contact upon reasonable request.

## ACKNOWLEDGEMENTS

This project was funded by grants from the European Research Council (ERC StG MEMCIRCUIT, GA 758032) and German Research Foundation (DFG JA 1999/3-1) to S.N.J.

## AUTHOR CONTRIBUTIONS

A.R., D.H. and S.N.J. designed the experiments. A.R. and D.H. performed the experiments.

A.R. analyzed the data. A.R. and S.N.J. wrote the paper.

## REFERENCES

1. O’Connor, D.H., Huber, D., and Svoboda, K. (2009). Reverse engineering the mouse brain. Nature 461, 923–929. 10.1038/nature08539.

2. Gomez-Marin, A., Paton, J.J., Kampff, A.R., Costa, R.M., and Mainen, Z.F. (2014). Big behavioral data: psychology, ethology and the foundations of neuroscience. Nat Neurosci 17, 1455–1462. 10.1038/nn.3812.

3. Luo, L., Callaway, E.M., and Svoboda, K. (2018). Genetic Dissection of Neural Circuits: A Decade of Progress. Neuron 98, 865. 10.1016/j.neuron.2018.05.004.

4. Bussey, T.J., Holmes, A., Lyon, L., Mar, A.C., McAllister, K.A., Nithianantharajah, J., Oomen, C.A., and Saksida, L.M. (2012). New translational assays for preclinical modelling of cognition in schizophrenia: the touchscreen testing method for mice and rats. Neuropharmacology 62, 1191–1203. 10.1016/j.neuropharm.2011.04.011.

5. Nithianantharajah, J., McKechanie, A.G., Stewart, T.J., Johnstone, M., Blackwood, D.H., St Clair, D., Grant, S.G., Bussey, T.J., and Saksida, L.M. (2015). Bridging the translational divide: identical cognitive touchscreen testing in mice and humans carrying mutations in a disease-relevant homologous gene. Sci Rep 5, 14613. 10.1038/srep14613.

6. Beck, J.A., Lloyd, S., Hafezparast, M., Lennon-Pierce, M., Eppig, J.T., Festing, M.F., and Fisher, E.M. (2000). Genealogies of mouse inbred strains. Nat Genet 24, 23–25. 10.1038/71641.

7. Huang, Z.J., and Zeng, H. (2013). Genetic approaches to neural circuits in the mouse. Annu Rev Neurosci 36, 183–215. 10.1146/annurev-neuro-062012-170307.

8. Harris, J.A., Hirokawa, K.E., Sorensen, S.A., Gu, H., Mills, M., Ng, L.L., Bohn, P., Mortrud, M., Ouellette, B., Kidney, J., et al. (2014). Anatomical characterization of Cre driver mice for neural circuit mapping and manipulation. Front Neural Circuits 8, 76. 10.3389/fncir.2014.00076.

9. Madisen, L., Garner, A.R., Shimaoka, D., Chuong, A.S., Klapoetke, N.C., Li, L., van der Bourg, A., Niino, Y., Egolf, L., Monetti, C., et al. (2015). Transgenic mice for intersectional targeting of neural sensors and effectors with high specificity and performance. Neuron 85, 942–958. 10.1016/j.neuron.2015.02.022.

10. Oh, S.W., Harris, J.A., Ng, L., Winslow, B., Cain, N., Mihalas, S., Wang, Q., Lau, C., Kuan, L., Henry, A.M., et al. (2014). A mesoscale connectome of the mouse brain. Nature 508, 207–214. 10.1038/nature13186.

11. Zhang, T., Zeng, Y., and Xu, B. (2017). A computational approach towards the microscale mouse brain connectome from the mesoscale. J Integr Neurosci 16, 291–306. 10.3233/JIN-170019.

12. Zingg, B., Hintiryan, H., Gou, L., Song, M.Y., Bay, M., Bienkowski, M.S., Foster, N.N., Yamashita, S., Bowman, I., Toga, A.W., and Dong, H.W. (2014). Neural networks of the mouse neocortex. Cell 156, 1096–1111. 10.1016/j.cell.2014.02.023.

13. Burgess, C.P., Lak, A., Steinmetz, N.A., Zatka-Haas, P., Bai Reddy, C., Jacobs, E.A.K., Linden, J.F., Paton, J.J., Ranson, A., Schroder, S., et al. (2017). High-Yield Methods for Accurate Two-Alternative Visual Psychophysics in Head-Fixed Mice. Cell Rep 20, 2513–2524. 10.1016/j.celrep.2017.08.047.

14. Andermann, M.L., Kerlin, A.M., and Reid, R.C. (2010). Chronic cellular imaging of mouse visual cortex during operant behavior and passive viewing. Front Cell Neurosci 4, 3. 10.3389/fncel.2010.00003.

15. Sanders, J.I., and Kepecs, A. (2012). Choice ball: a response interface for two-choice psychometric discrimination in head-fixed mice. J Neurophysiol 108, 3416–3423. 10.1152/jn.00669.2012.

16. Hangya, B., Ranade, S.P., Lorenc, M., and Kepecs, A. (2015). Central Cholinergic Neurons Are Rapidly Recruited by Reinforcement Feedback. Cell 162, 1155–1168. 10.1016/j.cell.2015.07.057.

17. Guo, Z.V., Li, N., Huber, D., Ophir, E., Gutnisky, D., Ting, J.T., Feng, G., and Svoboda, K. (2014). Flow of cortical activity underlying a tactile decision in mice. Neuron 81, 179–194. 10.1016/j.neuron.2013.10.020.

18. Abraham, N.M., Guerin, D., Bhaukaurally, K., and Carleton, A. (2012). Similar odor discrimination behavior in head-restrained and freely moving mice. PLoS One 7, e51789. 10.1371/journal.pone.0051789.

19. Resulaj, A., and Rinberg, D. (2015). Novel Behavioral Paradigm Reveals Lower Temporal Limits on Mouse Olfactory Decisions. J Neurosci 35, 11667–11673. 10.1523/JNEUROSCI.4693-14.2015.

20. Liu, D., Gu, X., Zhu, J., Zhang, X., Han, Z., Yan, W., Cheng, Q., Hao, J., Fan, H., Hou, R., et al. (2014). Medial prefrontal activity during delay period contributes to learning of a working memory task. Science 346, 458–463. 10.1126/science.1256573.

21. Thorndike, E.L. (1911). Animal intelligence; experimental studies (The Macmillan Company).

22. Skinner, B.F. (1938). The behavior of organisms: An experimental analysis (BF Skinner Foundation).

23. Sutton, R.S., and Barto, A.G. (2018). Reinforcement Learning, second edition: An Introduction (MIT Press).

24. Sanchez-Roige, S., Pena-Oliver, Y., and Stephens, D.N. (2012). Measuring impulsivity in mice: the five-choice serial reaction time task. Psychopharmacology (Berl) 219, 253–270. 10.1007/s00213-011-2560-5.

25. Sharma, S., Rakoczy, S., and Brown-Borg, H. (2010). Assessment of spatial memory in mice. Life Sci 87, 521–536. 10.1016/j.lfs.2010.09.004.

26. Lee, J.J., Krumin, M., Harris, K.D., and Carandini, M. (2022). Task specificity in mouse parietal cortex. Neuron 110, 2961–2969 e2965. 10.1016/j.neuron.2022.07.017.

27. Aoki, R., Tsubota, T., Goya, Y., and Benucci, A. (2017). An automated platform for high-throughput mouse behavior and physiology with voluntary head-fixation. Nat Commun 8, 1196. 10.1038/s41467-017-01371-0.

28. Lyamzin, D.R., Aoki, R., Abdolrahmani, M., and Benucci, A. (2021). Probabilistic discrimination of relative stimulus features in mice. Proc Natl Acad Sci U S A 118. 10.1073/pnas.2103952118.

29. Xiao, D., and Balbi, M. (2025). Continuous Auditory Feedback Promotes Fine Motor Skill Learning in Mice. eNeuro 12. 10.1523/ENEURO.0008-25.2025.

30. International Brain, L., Aguillon-Rodriguez, V., Angelaki, D., Bayer, H., Bonacchi, N., Carandini, M., Cazettes, F., Chapuis, G., Churchland, A.K., Dan, Y., et al. (2021). Standardized and reproducible measurement of decision-making in mice. Elife 10. 10.7554/eLife.63711.

31. Selen, L.P., Shadlen, M.N., and Wolpert, D.M. (2012). Deliberation in the motor system: reflex gains track evolving evidence leading to a decision. J Neurosci 32, 2276–2286. 10.1523/JNEUROSCI.5273-11.2012.

32. Gold, J.I., and Shadlen, M.N. (2001). Neural computations that underlie decisions about sensory stimuli. Trends Cogn Sci 5, 10–16. 10.1016/s1364-6613(00)01567-9.

33. Resulaj, A., Kiani, R., Wolpert, D.M., and Shadlen, M.N. (2009). Changes of mind in decision-making. Nature 461, 263–266. 10.1038/nature08275.

34. Kawato, M. (1999). Internal models for motor control and trajectory planning. Current Opinion in Neurobiology.

35. Mazzoni, P., and Krakauer, J.W. (2006). An implicit plan overrides an explicit strategy during visuomotor adaptation. J Neurosci 26, 3642–3645. 10.1523/JNEUROSCI.5317-05.2006.

36. Coen, P., Sit, T.P.H., Wells, M.J., Carandini, M., and Harris, K.D. (2023). Mouse frontal cortex mediates additive multisensory decisions. Neuron. 10.1016/j.neuron.2023.05.008.

37. Dadarlat, M.C., O’Doherty, J.E., and Sabes, P.N. (2015). A learning-based approach to artificial sensory feedback leads to optimal integration. Nat Neurosci 18, 138–144. 10.1038/nn.3883.

38. Flesher, S.N., Downey, J.E., Weiss, J.M., Hughes, C.L., Herrera, A.J., Tyler-Kabara, E.C., Boninger, M.L., Collinger, J.L., and Gaunt, R.A. (2021). A brain-computer interface that evokes tactile sensations improves robotic arm control. Science 372, 831–836. 10.1126/science.abd0380.

39. Attinger, A., Wang, B., and Keller, G.B. (2017). Visuomotor Coupling Shapes the Functional Development of Mouse Visual Cortex. Cell 169, 1291–1302 e1214. 10.1016/j.cell.2017.05.023.

40. Meijer, G.T., Pie, J.L., Dolman, T.L., Pennartz, C.M.A., and Lansink, C.S. (2018). Audiovisual Integration Enhances Stimulus Detection Performance in Mice. Front Behav Neurosci 12, 231. 10.3389/fnbeh.2018.00231.

41. Siemann, J.K., Muller, C.L., Bamberger, G., Allison, J.D., Veenstra-VanderWeele, J., and Wallace, M.T. (2015). A novel behavioral paradigm to assess multisensory processing in mice. Frontiers in Behavioral Neuroscience 8. 10.3389/fnbeh.2014.00456.

42. Burke, J.L., Prewett, M.S., Gray, A.A., Yang, L., Stilson, F.R.B., Coovert, M.D., Elliot, L.R., and Redden, E. (2006). Comparing the effects of visual-auditory and visual-tactile feedback on user performance: a meta-analysis. Proceedings of the 8th international conference on Multimodal interfaces. Association for Computing Machinery.

43. Sigrist, R., Rauter, G., Riener, R., and Wolf, P. (2013). Augmented visual, auditory, haptic, and multimodal feedback in motor learning: a review. Psychon Bull Rev 20, 21–53. 10.3758/s13423-012-0333-8.

44. Hwang, J., Mitz, A.R., and Murray, E.A. (2019). NIMH MonkeyLogic: Behavioral control and data acquisition in MATLAB. J Neurosci Methods 323, 13–21. 10.1016/j.jneumeth.2019.05.002.

45. Heffner, H.E., and Heffner, R.S. (2007). Hearing ranges of laboratory animals. J Am Assoc Lab Anim Sci 46, 20–22.

46. Radziwon, K.E., and Dent, M.L. (2014). Frequency difference limens and auditory cue trading in CBA/CaJ mice (Mus musculus). Behav Processes 106, 74–76. 10.1016/j.beproc.2014.04.016.

47. Radziwon, K.E., June, K.M., Stolzberg, D.J., Xu-Friedman, M.A., Salvi, R.J., and Dent, M.L. (2009). Behaviorally measured audiograms and gap detection thresholds in CBA/CaJ mice. J Comp Physiol A Neuroethol Sens Neural Behav Physiol 195, 961–969. 10.1007/s00359-009-0472-1.

48. Schmitt, L.I., Wimmer, R.D., Nakajima, M., Happ, M., Mofakham, S., and Halassa, M.M. (2017). Thalamic amplification of cortical connectivity sustains attentional control. Nature 545, 219–223. 10.1038/nature22073.

49. Wimmer, R.D., Schmitt, L.I., Davidson, T.J., Nakajima, M., Deisseroth, K., and Halassa, M.M. (2015). Thalamic control of sensory selection in divided attention. Nature 526, 705–709. 10.1038/nature15398.

